# Surface-Enhanced Raman Spectroscopy-Assisted Lateral Flow Test for Adenine and IgG Analysis

**DOI:** 10.1101/2025.06.11.659022

**Authors:** Zuzana Chaloupková, Klára Gajdošová, Kateřina Poláková, Václav Ranc

## Abstract

The development of analytical methods allowing a fast and easy to perform chemical analysis of complex samples under non-laboratory conditions presents a considerable scientific challenge that requires attention. Many analytical tasks are performed using commercially available immunochemistry based lateral flow tests, where the results are detected via a direct observation of the color change of the test strip. However, these tests in many cases do not have the desired levels of selectivity and/or sensitivity or even in some cases introduce an unwanted degree of subjectivity and thus uncertainty. Here, we developed a novel lateral flow analysis test designed for easy-to-run and easily to-interpret detection implementing surface enhanced Raman scattering. The strip utilizes modular polymeric sections specifically engineered to simplify the separation of target analytes from complex biological samples. This separation relies on a series of carefully designed physicochemical interactions between the sample components and the polymeric materials embedded in the strip. A key feature of this detection method is an analytical area on the strip that incorporates plasmonic silver nanostructures with a possibility of further surface functionalization using various selectors. We evaluated this test on the selective detection of IgG through its antibody and Adenine, as representatives of proteins and low molecular compounds. This innovative test format combines modular design with advanced detection, offering significant potential for applications in biomedical research, diagnostic testing, and monitoring of immune function.

**Highlights:**

- Reliable and fast detection of low-molecular-weight substances and proteins.
- Combination of modular polymer lateral flow tests and SERS analysis.
- Detection of complex samples, pre-separation, and pre-concentration of analytes prior to the detection.

**Graphical Abstract:** 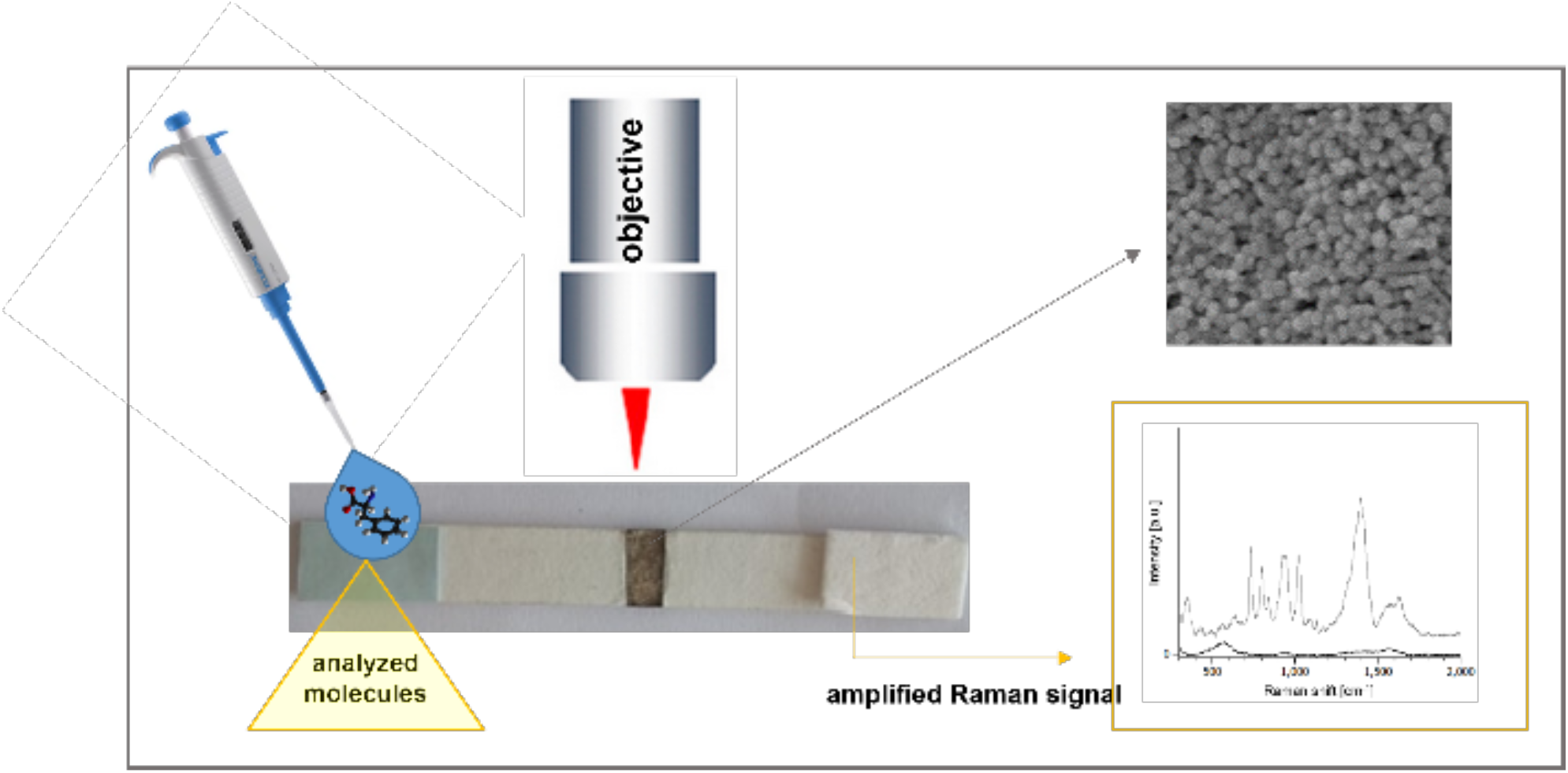

## 1 Introduction

The demand for suitable sensitive and selective analytical techniques arises from the need to accurately detect and quantify trace amounts of substances in complex matrices, including pharmaceutical, clinical, environmental or forensic samples [1]. Currently, applied techniques such as chromatography and mass spectrometry, though highly effective, often require extensive sample preparation, longer analysis times, and importantly also higher running costs. Raman spectroscopy offers a promising alternative due to its ability to provide rapid, non-destructive, and highly specific molecular fingerprinting without the need for extensive sample preparation [2]. A notable advancement in this field is Surface-Enhanced Raman Scattering (SERS), which significantly improves detection limits by utilizing nanomaterials with specific optoelectronic properties, particularly those exhibiting strong surface plasmon resonance. While traditional SERS substrates have relied heavily on colloidal silver or gold nanoparticles, these systems present challenges related to stability, reproducibility, and inhomogeneity due to varying particle sizes and concentrations. Alternatively, solid substrates like metal films or glass coated with gold or silver nanoparticles offer enhanced performance, although their preparation can be labor-intensive and costly, alongside issues of fragility. Together, these aspects highlight the intricacies and advancements in Raman spectroscopy, positioning it as a vital tool for molecular analysis [3, 4].

Recent advances have integrated surface-enhanced Raman scattering (SERS) with lateral flow assays (LFA) to increase the sensitivity and specificity of point-of-care diagnostics. These SERS-based LFAs often use electrostatic interactions to attach detection probes to the test membrane. For example, Hwang et al. developed a SERS-based LFA for the detection of staphylococcal enterotoxin B (SEB) [5]. They used hollow gold nanospheres labeled with Raman reporters as detection probes that electrostatically interacted with the test membrane to identify SEB antigens. This approach allowed rapid identification by color change and quantitative evaluation by Raman signal measurement. Similarly, Chen et al. [6] introduced a SERS-based LFA strip for multiplexed detection of anti-SARS-CoV-2, IgM, and IgG antibodies. They used gold-silver core-shell nanostructures as SERS probes, which adhered to the test membrane through electrostatic interactions. This construct achieved detection limits significantly lower than traditional colorimetric methods, increasing sensitivity for diagnosing infection at an early stage. However, relying solely on electrostatic interactions to attach probes can lead to challenges with stability, reproducibility, and homogeneity due to differences in particle size and concentration. Electrostatic binding is often sensitive to environmental changes that can affect the consistency and efficiency of the assay [7]. Covalent bonding methods have been proposed to address these problems. Covalent immobilization offers a stronger and more stable attachment of detection probes to the membrane, potentially improving the reliability and performance of SERS-based LFA. Incorporation of covalent bonding techniques could increase the robustness of SERS-based lateral flow assays, ensure consistent probe attachment, and improve overall assay sensitivity and specificity [8].

Here we present a novel lateral-flow-based approach, which according to our best knowledge for the first time utilizes well-controlled covalent binding of silver nanostructures to the nitrocellulose membrane via nitrogen-containing functional groups. This functionalization method achieves homogeneity and considerably lowers detection limits down to the pg/L concentration levels, resulting in a stable analytical signal that outperforms traditional techniques that rely on electrostatic interactions. In addition, the robust anchoring of the nanoparticles prevents washout, increasing the reliability of the results. The developed analytical system was tested on the analysis of Adenine, as a model of low-molecular compound, and IgG to test the selective detection of biomacromolecules in complex samples.

## 2 Methodology Materials and Methods

### 2.1 Chemicals

Sterile Whatman membrane filter (0.45 µm pore size, 47 mm diameter, Sigma Aldrich, USA), Sodium tetrahydridoborate (powder, ≥98.0%, Sigma Aldrich, USA), Sodium citrate tribasic dihydrate (Molecular Biology Grade, Sigma Aldrich, USA), Sodium hydroxide (monohydrate, 99.99 Suprapur^®^, Sigma Aldrich, USA), adenine (powder, ≥99%, Sigma Aldrich, USA), Anti-Mouse IgG antibody (affinity isolated antibody, lyophilized powder, Sigma Aldrich, USA), IgG from mouse serum (reagent grade, ≥95% (SDS-PAGE), lyophilized powder, Sigma Aldrich, USA), BSA (lyophilized powder, ≥96%, Sigma Aldrich, USA), PSA (from human semen, Sigma Aldrich, USA).

### 2.2. Method of preparing the test strip

The test strips were prepared according to the procedure described in the European patent [9]. A nitrocellulose membrane with a pore size of 30 µm, specially selected for its filtration properties, forms the base of the strip. The membrane serves as a key component in the separation and detection process. At one end of the strip, referred to as the sample section, the nitrocellulose membrane is attached to a polyvinyl chloride plastic substrate to facilitate sample application without risk of leakage. The opposite end of the strip is designated as the absorption section, which uses a sterile Whatman membrane filter (0.45 µm pore size, 47 mm diameter, Sigma Aldrich, USA) from a commercial source to capture the analyte. Central to the functionality of the strip is part D (shown in Figure 1), where silver nanoparticles are covalently bonded to the nitrocellulose membrane. This section serves as a substrate for surface-enhanced Raman spectroscopy (SERS) and allows for the capture and subsequent detection of target compounds. The final dimensions of the prepared test strip are 70 mm x 20 mm x 0.5 mm; however, these dimensions are flexible and allow producing strips of different sizes tailored to specific separation needs. For the sampling section, a Whatman membrane filter with a pore size of 0.30 µm was used, while the absorption section used a sterile membrane filter as mentioned above.

**Fig. 1:**
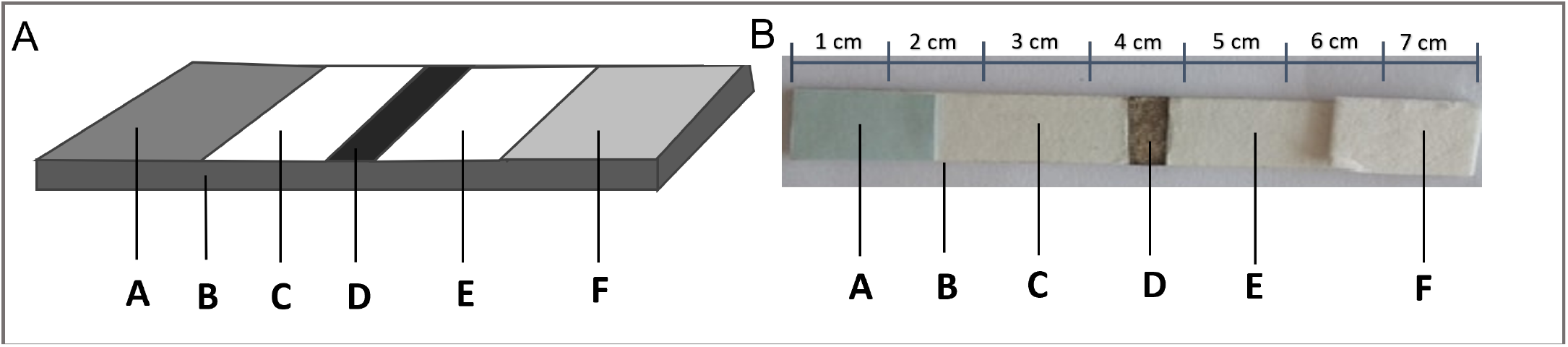
A: shows a schematic drawing of a test strip containing silver nanoparticles as a substrate for surface-enhanced Raman spectroscopy. B: photograph of strip, with scale.

### 2.3. Method for the preparation of silver nanoparticles covalently bonded to a nitrocellulose membrane

Silver nanoparticles were prepared according to the described protocol [10] using NaBH4 as the primary reducing agent and trisodium citrate (TSC) as the secondary reducing agent as well as stabilizer. A freshly prepared 45 mL aqueous solution containing 50 mmol/L NaBH4 and 2 mmol/L TSC was stirred for 30 min at 60 °C in the dark. A 5 ml solution of silver nitrate at a concentration of 1,22 mmol/L was then added dropwise to the solution, the temperature was subsequently raised to 90 °C and the pH of the reaction mixture was adjusted to 10,5 with 0,1 M NaOH. The reaction was run for 20 min until there was no visible color change in the reaction mixture. The mixture was then allowed to cool to room temperature and the silver nanoparticles were separated by centrifugation (12,000 rpm, 15 minutes) and washed three times with water. Their immobilization on nitrocellulose membrane was performed by inserting a nitrocellulose membrane (substrate) of l × l cm and 30 µm pore size into a 10 mL tube. Further, 0.02 g of TSC dissolved in water with a final concentration of 35 mmol/L and silver nanoparticles prepared according to the above protocol in a colloidal suspension with a silver nanoparticle concentration of 9.01 ×10^11^ per mL in a final volume of 3 mL were added to the tube thus prepared. The mixture thus prepared was subjected to reduction for seven days under light-free conditions at room temperature. After seven days, the substrate was removed from the tube, rinsed several times with distilled water and allowed to dry.

### 2.4. Method for the preparation of silver nanoparticles electrostatically bonded to a nitrocellulose membrane

Silver nanoparticles were prepared according to the protocol previously described by Agnihotri [10] and the procedure described in the Section 2.3 Their immobilization on the nitrocellulose membrane is based on the electrostatic binding of silver nanoparticles by dropping these particles in a volume of 50 µL (in a colloidal suspension with silver nanoparticles at a concentration of 9.01 x10^11^ per mL) onto a nitrocellulose membrane with a size of 1 × 1 cm and a pore size of 30 µm. To increase the number of immobilized particles, this step of dripping 50 µL of colloidal suspension was repeated 5 times, always after a complete drying. Once the substrate was completely dried out, it was cut to the size required for application to the test strip described in Section 2.2. The nitrocellulose membrane containing electrostatically anchored silver nanoparticles was attached to the plastic substrate using an adhesive.

### 2.5. Measurement Adenine using covalently bound silver nanoparticles

A test strip containing silver nanoparticles covalently bonded to a nitrocellulose membrane was used for measurements using surface-enhanced Raman spectroscopy. The test strip was tested for the detection of Adenine at a concentration 10 µg/mL to 100 pg/mL. The plasmonic strip was incubated in 200 µL of adenine solution for 30 mins. Measurements on the Raman microscope were performed after the entire test strip had dried. An intelligent depolarized high-brightness laser, solid-state, diode-pumped with constant power, with an excitation wavelength of 532 nm and laser power on the sample of 0.1 mW, 32 exposures per spectrum, was used for excitation.

### 2.6. Preparation of silver nanoparticles with immobilized anti-IgG antibody by direct adsorption as a substrate for IgG detection

A 1 × 1 cm nitrocellulose membrane strip with a pore size of 30 µm was immersed in 500 µL of aqueous solution containing 1 ng/L anti-IgG antibody. A total of 3 mL of colloidal silver nanoparticles prepared according to the procedure described in section 2.3 (with a concentration of 9.01 × 10^11^ silver nanoparticles per mL) were then added to the reaction mixture. This mixture was then shaken for 45 minutes to facilitate binding. After the incubation period, the nitrocellulose membrane was allowed to dry, resulting in the binding of the anti-IgG antibody to the membrane. The dried membrane, now containing the bound anti-IgG antibody, was transferred to a 10 mL tube. Then 0.02 g sodium borohydride (NaBH4) was added to give a total concentration of 30 mM, together with the previously prepared silver nanoparticles containing the anti-IgG antibody, adjusting the final volume to 3 mL. The resulting substrate was subjected to a five-day reduction process, which was maintained in the absence of light at room temperature. At the end of this period, the substrate was carefully removed from the tubes, rinsed several times with distilled water to remove any unbound material, and then allowed to dry completely. The final product was a nitrocellulose membrane with silver nanoparticles that had anti-IgG antibodies covalently bound to their surface, as well as additional anti-IgG antibodies immobilized directly on the nitrocellulose membrane itself. After complete drying, the membrane was cut to the required dimensions for incorporation into the test strip.

### 2.7. Preparation of nitrocellulose membrane with electrostatically bound silver nanoparticles and immobilized anti-IgG antibody

Immobilization of the anti-IgG antibody was performed by direct adsorption to the nitrocellulose membrane. A 1 × 1 cm membrane with electrostatically bound silver nanoparticles was immersed at room temperature for 45 min in 500 µL of an aqueous solution of the appropriate antibody (anti-IgG, c = 1 ng/L). The result was a nitrocellulose membrane to which the silver nanoparticles and immobilized anti-IgG antibody were electrostatically bound. After complete drying, the membrane was cut to the required dimensions for incorporation into the test strip.

### 2.8. Analysis of IgG

The functionalized test strip was applied in the detection of IgG protein at a concentration of 1 ng/mL using functionalized silver nanoparticles. This analytics was achieved through direct adsorption and electrostatic interactions with the antibodies on the surface of the nanoparticles. A 50 µL sample of the 1 ng/mL IgG solution was applied to the test strip, which included Part D featuring covalently bonded silver nanoparticles and immobilized anti-IgG antibodies, as outlined in Sections 2.6 and 2.7. After the sample had fully traversed the strip and dried, Raman measurements were performed. The excitation was provided by a laser with a wavelength of 532 nm, delivering a power of 0.1 mW per sample. This method enabled sensitive detection of the IgG protein through Raman spectroscopy.

### 2.9. Analysis of mixture containing IgG, PSA and BSA

The test strip was evaluated for the detection of IgG protein at a concentration of 1 ng/ml in a mixture containing IgG, prostate specific antigen (PSA) and bovine serum albumin (BSA) using functionalized silver nanoparticles. This was achieved by direct adsorption of the antibody onto the surface of the silver nanoparticles. A 50 µL sample of the mixture, which had a total protein concentration of 1 ng/mL with a 1:1:1 (IgG : PSA : BSA) ratio, was applied to the test strip (Part A). This strip contained part D, which contained covalently bound silver nanoparticles and immobilized anti-IgG antibodies as described in Section 2.5. The excitation source was a He-Ne laser with a wavelength of 532 nm, which delivered a power of 0.1 mW per sample, allowing sensitive detection of the IgG protein by Raman spectroscopy.

### 2.10. Apparatus

Raman spectra were measured on a DXR Raman spectroscope (Thermo Scientific, U.S.A) equipped with a red excitation laser operating at 532 nm. Stokes Raman spectra were collected in the 500-2000 cm^−1^ range with a spectral resolution of 1.0 cm^−1^. The Raman spectrometer was operated with the following experimental parameters: exposure time 32 s, laser power on sample 0.1 mW, 32 micro-scans were averaged to obtain one experimental point. Spectral background was corrected using subtraction of polynomic functions (n = 2) to remove interference caused by fluorescence. The measured spectra are presented in terms of their Raman scattered photon count. The samples were analyzed by Scanning Electron Microscope (SEM) Hitachi SU6600 with accelerating voltage 5 kV.

## 3 Results and discussion

The lateral flow test is based on the covalent anchoring of silver nanoparticles on a nitrocellulose substrate, which is consequently incorporated into the strip structure. The test is also equipped with a substrate that prevents the flow of liquid through the porous nitrocellulose membrane outside of the strip. Attached to this membrane is an additional nitrocellulose layer with a pore size of 0.30 µm, on which silver nanoparticles are covalently anchored. According to dynamic light scattering (DLS) analysis, the silver nanoparticles used in this study exhibit a narrow size distribution. Based on the findings of Agnihotri et al., the average hydrodynamic diameter of citrate-stabilized silver nanoparticles was reported as 43.2 ± 3.1 nm, indicating a relatively monodisperse population. These values correspond to the typical size range used in lateral flow tests, ensuring optimal mobility through the porous nitrocellulose membrane and efficient generation of a visual signal. The strip includes a sample application section with the same pore size of nitrocellulose and is modified for adhesion with a non-wetting surface, allowing for a large capture volume more than 50 µl/cm^2^. At the opposite end of the strip is the absorption section, where the nitrocellulose membrane has a larger pore size of 0.45 to 0.50 µm, facilitating efficient sample absorption. The configuration of the strip allows seamless transfer of the liquid sample from the application section through the nitrocellulose membrane and nanoparticles to the absorption section, ensuring efficient detection.

The test strip for substance detection works on a simple but effective mechanism (Figure 1). First, a liquid sample of a mixture containing the substance of interest is applied to the sample area (A) of the strip, it gradually rises through the nitrocellulose membrane (C). The liquid then encounters a portion (D) containing silver nanoparticles, which plays a key role in the detection process. As the sample moves, the target substance adsorbs onto the surface of present plasmonic nanoparticles. This adsorption separates the target substance from the rest of the mixture and concentrates it at that specific location. The presence of the nanoparticles enhances the Raman signal of the adsorbed substance through the mechanism of surface-enhanced Raman spectroscopy (SERS), allowing for sensitive detection. Finally, the sample advances to the region (E) where any remaining liquid is captured on the absorption part (F). This design ensures accurate identification and quantification of the target substance based on the amplified Raman signal generated by interaction with the nanoparticles.

Subsequently, the strip was characterized by SEM (Figure 2). The figure shows the nitrocellulose membrane at the strip. Shows uniform coverage of the nitrocellulose membrane with silver nanoparticles. Confirmation of Ag-N binding is shown in the Raman spectrum (Fig. 2D).

**Fig. 2.**
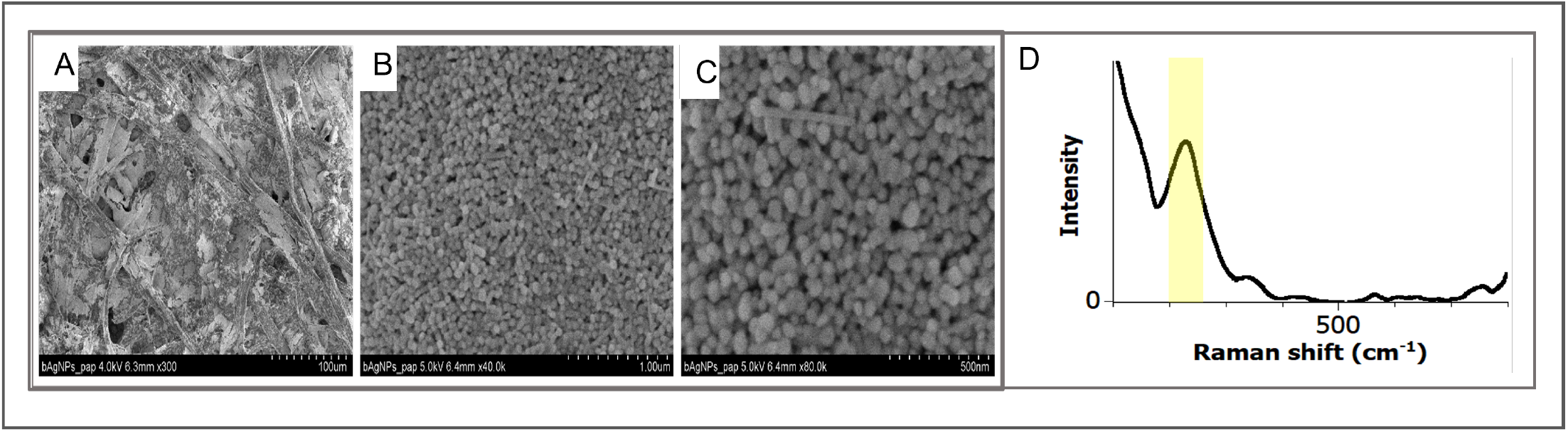
shows an electron microscope image showing silver nanoparticles (light spherical objects) covalently immobilized on nitrocellulose membrane (dark background). A) Magnifications of 100 µm, B) 1 µm, C) 500 nm. D) The bond observed in the Raman spectrum around 220 cm^-1^ corresponding to the vibrational mode of the Ag-N bond.

Confirmation of Ag–N (silver–nitrogen) bonding is a critical step in verifying the successful functionalization or interaction of silver nanoparticles (AgNPs) with biomolecules or ligands containing nitrogen groups [11]. The presence of Ag–N bonds indicates that the nanoparticles are effectively integrated. This integration is crucial for specific molecular recognition and signal generation in biosensor applications. Silver nanoparticles are also known to enhance Raman signals through surface-enhanced Raman scattering (SERS). Ag–N interactions can influence the local electronic environment and surface chemistry of nanoparticles, contributing to stronger and more reliable Raman signals. Confirmation of the Ag–N bond provides evidence that silver nanoparticles are not only present on the membrane but are also tightly bound. This considerably increases the sensitivity of the detection system, especially in bioanalytical or diagnostic applications [12].

The sensing capabilities of covalently immobilized silver nanoparticles (AgNPs) on nitrocellulose test strips were deeply evaluated using adenine as a low molecular weight model Raman probe molecule. The reference Raman spectrum of adenine at a concentration of 1 µg/ml showed a characteristic vibration band at 732 cm−^1^, which was assigned to ring breathing modes (Figure 3A), in agreement with previous literature [13, 14]. This band served as the main analytical marker in subsequent SERS measurements. Comparison of adenine detection using colloidal AgNPs and AgNPs covalently bound to strips revealed a significantly increased SERS response for the latter (Figure 3B). The intensity of the adenine band at 732 cm−^1^ was significantly higher on the AgNPs@strips platform, with an enhancement factor (EF) of 3.5 × 10^3^, compared to only 4.9 × 10^2^ for colloidal AgNPs. This increase of almost one order of magnitude highlights the importance of immobilizing nanoparticles via Ag–N covalent interactions, which ensure stable and homogeneous distribution of plasmonic hotspots. Similar immobilization strategies have demonstrated improved sensitivity and reproducibility in other SERS-based biosensors [15, 16, 17].

**Figure 3.**
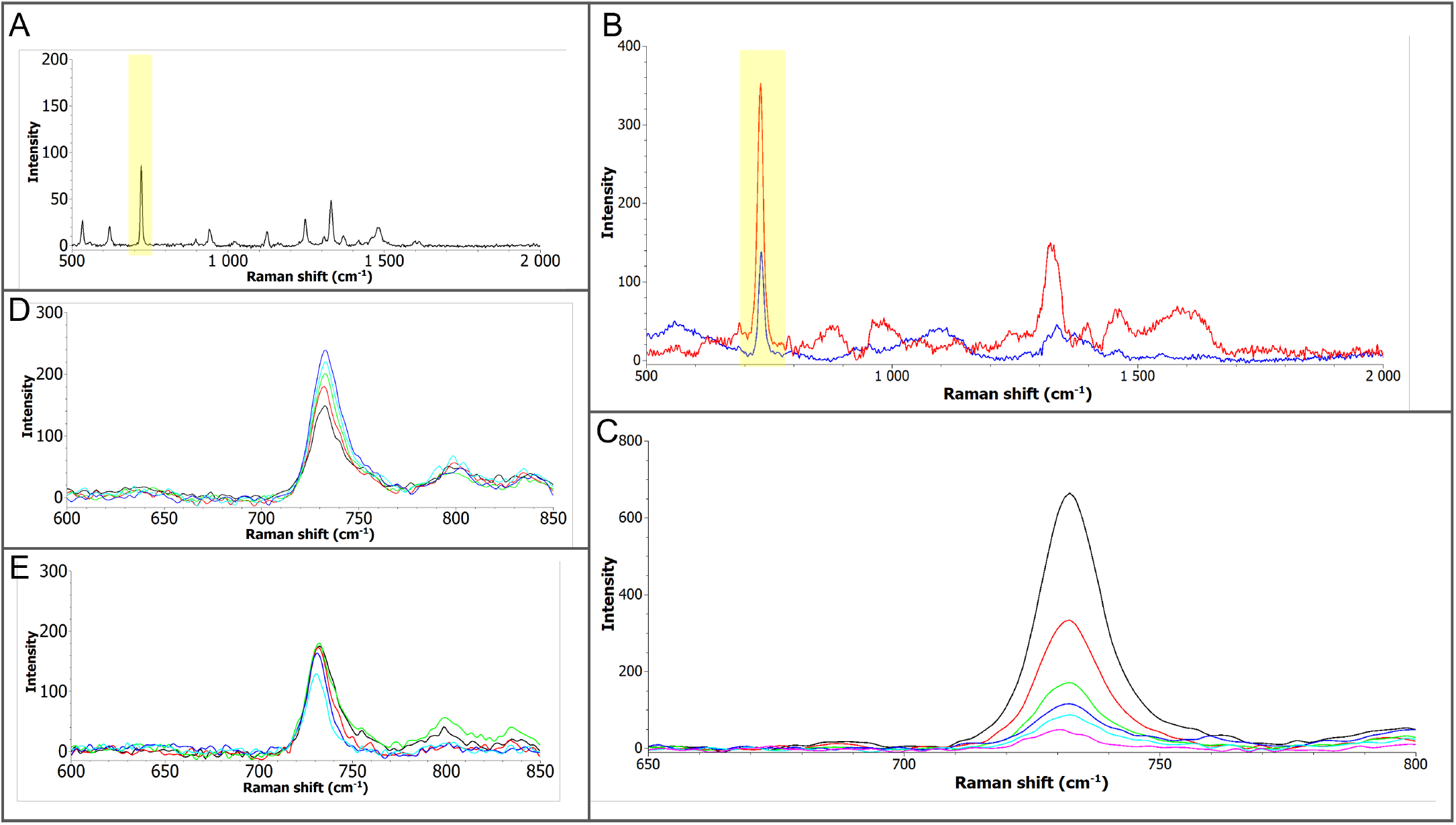
A) Raman spectrum of an Adenine standard (c = 1 mg/mL) with the typical Adenine band in the 732 cm^-1^ region (ring stretching) highlighted in yellow; B) Raman spectrum of Adenine (c = 1 µg/mL) measured on colloidal silver nanoparticles (blue) and on a strip (red), with the typical Adenine band in the 732 cm-1 region highlighted in yellow; C) SERS intensity dependence for the 732 cm^-1^ band on Adenine concentration measured on strips (10 µg/mL – dark blue, 1 µg/mL - blue, 100 ng/mL - green, 10 ng/mL - red and 10 pg/mL - black); D) Repeatability of measurements on one strip at five different spots (Adenine band 732 cm^-1^, c = 1 ng/mL); E) Repeatability of measurements on five different strips (Adenine band 732 cm^-1^, c = 1 ng/mL).

The developed SERS platform demonstrated impressive analytical performance and achieved a limit of detection (LOD) for adenine of only 4 pg/ml (Figure 3C). To our knowledge, this LOD surpasses many previously reported paper- or strip-based SERS systems, which typically achieve LODs in the low nanogram to sub-nanogram range [18, 19]. Furthermore, the linear dynamic range covers five orders of magnitude (1000 ng/ml to 0.01 ng/ml) with a correlation coefficient R^2^=0.81, indicating the quantitative potential of the platform. The robustness of the manufactured strips was further verified by evaluating the repeatability both between five spatially distinct points on a single strip (RSD = 9.9%) and between five independently prepared strips (RSD = 3.4%) at an adenine concentration of 1 ng/ml (Figures 3D and 3E). These low relative standard deviations confirm the reproducibility of the manufacturing process and further support the uniform distribution and stable binding of AgNP to the nitrocellulose membrane.

Together, these results demonstrate that covalently immobilized AgNPs significantly enhance the sensitivity and reproducibility of SERS on paper substrates. This approach holds promises for the development of cost-effective portable biosensors with ultrasensitive detection capabilities for small molecules.

Sensitive and selective detection of protein biomarkers is essential in diagnostic and analytical applications. To further evaluate an available selectivity of the system, the here developed substrates were applied in a detection of IgG in a mixture containing other relevant proteins. Figure 4A shows the Raman spectrum of a pure, unmodified strip, which exhibits negligible background signals. This confirms that the nitrocellulose substrate *per se* is inherently SERS-inactive and suitable for electrostatic-controlled detection of biomolecules. Figure 4B shows the SERS spectrum of anti-IgG antibodies (c = 1 ng/mL) immobilized on the AgNP-coated surface. Although the signal is relatively weak, newly emerging spectral characteristics indicate successful attachment of antibodies to the surface via electrostatic interactions, which serve as recognition elements for subsequent IgG detection. Figure 4C shows the SERS IgG (1 ng/mL) spectrum obtained after applying IgG (1 ng/mL) to the functionalized strip. Distinct vibration bands are clearly visible, especially in the region between 1000–1700 cm−^1^ (highlighted in yellow). These bands are associated with protein-specific modes such as amide I, amide III, and side chains of aromatic amino acids such as phenylalanine and tryptophan. The clear spectral fingerprint confirms that AgNPs effectively amplify Raman signals even for proteins weakly bound by electrostatic attraction, confirming the sensitivity of this approach [20]. Figures 4D and 4E show the spectra of BSA (bovine serum albumin) and PSA (prostate-specific antigen) at the same concentration (1 ng/ml). These proteins produce unique spectral profiles that differ from IgG, demonstrating the strip’s potential for differential protein recognition based on their intrinsic Raman vibration modes, even without covalent or site-specific binding.

**Figure 4.**
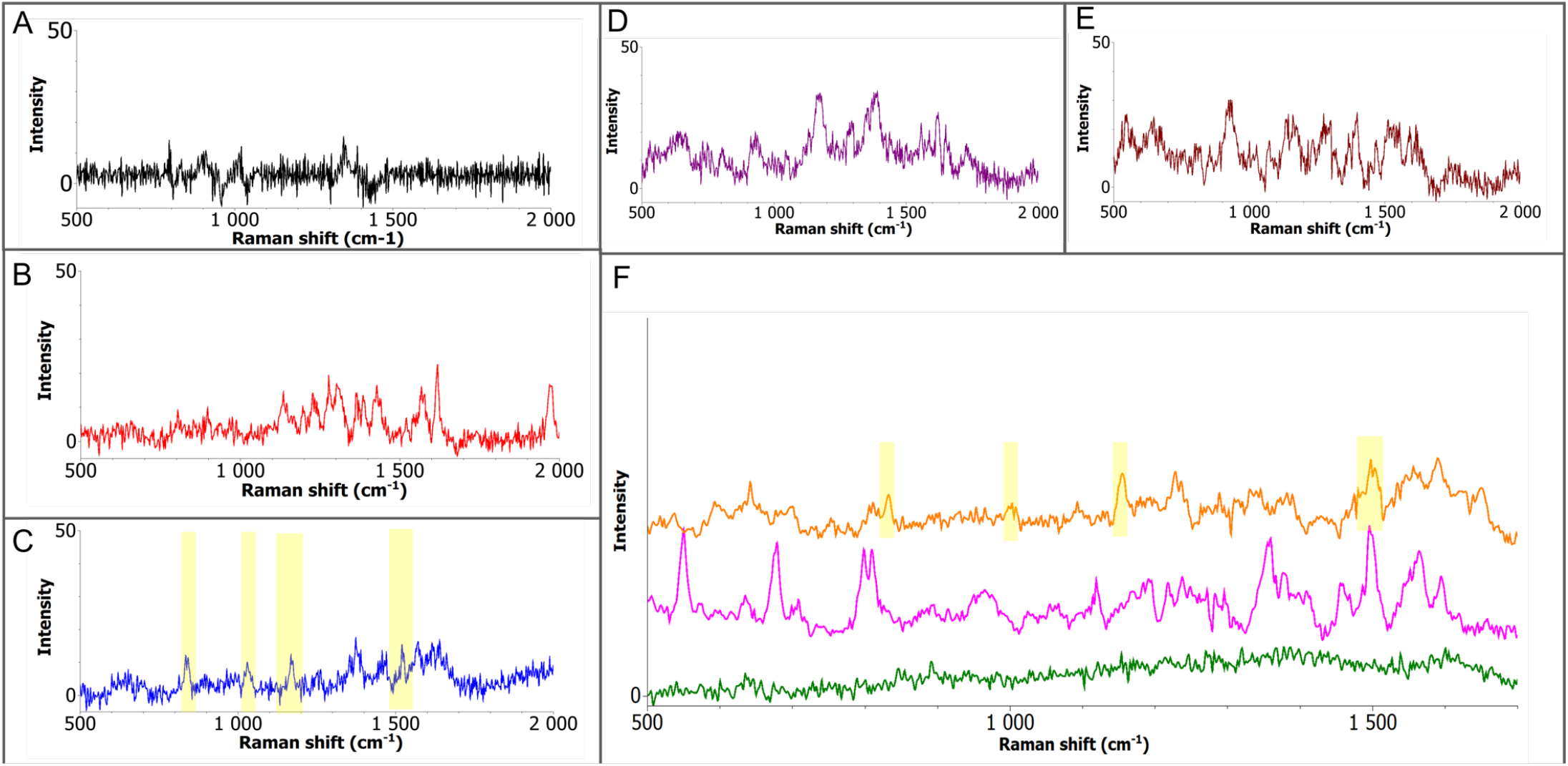
A) Raman spectrum of pure strips; B) Raman spectrum of anti-IgG measured on strips (c=1 ng/mL); C) Raman spectrum of IgG measured on strips (c=1 ng/mL) with IgG bands highlighted in yellow; D) Raman spectrum of BSA measured on strips (c=1 ng/mL); E) Raman spectrum of PSA measured on strips (c=1 ng/mL); F) Raman spectra of anti-IgG with IgG (where anti-IgG was bound electrostatically - green) or by direct absorption (pink) and Raman spectrum of anti-IgG with IgG in a mixture (containing IgG, BSA and PSA) with yellow bands for IgG, c=1 ng/mL

Figure 4F compares IgG detection in three different scenarios, where the green spectrum represents IgG detection on strips with immobilized anti-IgG. The signal is relatively low and close to the background, suggesting that the binding between anti-IgG and IgG alone is relatively weak and may lead to lower surface affinity or suboptimal orientation of IgG molecules. The pink spectrum shows a stronger SERS signal from IgG applied to strips with pre-immobilized anti-IgG. This improvement may be the result of increased local IgG concentration near the anti-IgG binding zones, allowing partial recognition based on affinity even under non-covalent conditions. The orange spectrum shows the detection of IgG in a complex mixture containing BSA and PSA. Despite the presence of potentially interfering proteins, the Raman bands specific for IgG remain visible, suggesting selective enrichment and detection, likely due to residual specificity retained by anti-IgG through weak interactions.

These results suggest that even without covalent or highly specific immobilization methods, covalently functionalized AgNP@strips can enable nanoparticle-enhanced SERS detection of target proteins at the nanogram per milliliter level. While signal intensity is generally lower than in systems using covalent bonding, the method retains clear protein discrimination capabilities and exhibits sufficient selectivity for use in simple biological matrices [15, 21]. Compared to more complex chemical surface modifications, direct adsorption represents a simple, inexpensive, and rapid approach to functionalizing SERS substrates. This makes the method attractive for the development of diagnostic platforms for point-of-care applications, especially in conditions where chemical modification or biological conjugation is not feasible.

## 4 Conclusions

In this study, we developed a new SERS-based lateral flow platform that utilizes the covalent immobilization of silver nanoparticles (AgNPs) on nitrocellulose membranes for highly sensitive and selective detection of both small molecules and protein biomarkers. In contrast to conventional paper-based SERS substrates, which mostly rely on electrostatic interactions, our approach introduces stable bonds between the immobilized silver nanoparticles and the substrate that also ensure uniform nanoparticle distribution, enhanced signal intensity and reproducibility, and significant resistance to washout or environmental variability.

The here developed test strips demonstrated an interesting analytical performance. For small molecule detection, the adenine model system achieved a limit of detection (LOD) of 4 pg/ml, surpassing most previously published paper- and strip-based SERS systems. Furthermore, a wide linear dynamic range and repeatability (RSD < 10%) confirmed the robustness and reproducibility of our manufacturing protocol. When used in protein analysis, AgNP@strips achieved sensitive and specific detection of IgG at 1 ng/ml even in complex mixtures containing potentially interfering proteins including PSA and BSA. Comparative experiments clearly showed that covalently immobilized AgNPs outperformed traditional electrostatic binding methods in terms of signal intensity, stability, and resolution.

Overall, our results highlight that covalent functionalization of AgNPs on paper substrates enables a powerful combination of SERS sensitivity with the simplicity of lateral flow formats. This hybrid platform overcomes many limitations of existing SERS-LFA systems and offers a low-cost, portable, and reliable solution for ultrasensitive detection in biomedical, environmental, and forensic applications. The versatility of this method makes it a promising candidate for future development of point-of-care diagnostics with multiplexing potential.

## Notes

The authors declare no conflict of interest.

